# IQD proteins integrate auxin and calcium signaling to regulate microtubule dynamics during *Arabidopsis* development

**DOI:** 10.1101/275560

**Authors:** Jos R. Wendrich, Bao-Jun Yang, Pieter Mijnhout, Hong-Wei Xue, Bert De Rybel, Dolf Weijers

## Abstract

Geometry and growth and division direction of individual cells are major contributors to plant organ shape and these processes are dependent on dynamics of microtubules (MT). Different MT structures, like the cortical microtubules, preprophase band and mitotic spindle, are characterized by diverse architectural dynamics (Hashimoto, 2015). While several MT binding proteins have been identified that have various effects on MT stability and architecture, they do not discriminate between the different MT structures. It is therefore likely that specific MT binding proteins exist that differentiate between MT structures in order to allow for the differences in architectural dynamics. Although evidence for the effect of specific cues, such as light and auxin, on MT dynamics has been shown in recent years (Lindeboom *et al.*, 2013; Chen *et al.*, 2014), it remains unknown how such cues are integrated and lead to specific effects. Here we provide evidence for how auxin and calcium signaling can be integrated to modulate MT dynamics, by means of IQD proteins. We show that the *Arabidopsis* IQD15-18 subclade of this family is regulated by auxin signaling, can bind calmodulins in a calcium-dependent manner and are evolutionarily conserved. Furthermore, AtIQD15-18 directly bind SPIRAL2 protein *in vitro* and *in vivo* and modulate its function, likely in a calmodulin-dependent way, thereby providing a missing link between two important regulatory pathways of MT dynamics.

**One sentence summary:** IQD proteins integrate auxin and calcium signaling, two major signaling pathways, to control the cytoskeleton dynamics and cell shape of *Arabidopsis*.

## Introduction

Plant organ shape is mainly controlled by the geometry of individual cells and the orientation of their cell divisions and growth axes. All these processes are highly dependent on microtubule (MT) dynamics that are part of the cytoskeleton of the cell (Lloyd and Chan, 2004). Disruption of these MT dynamics can lead to severe defects ranging from altered vesicle trafficking to misalignment of chromosomes during division and disordered division plane orientation (Kimata *et al.*, 2016). Furthermore, organization of the dynamic MTs at the cell cortex dictates the direction of cell expansion by guiding cellulose synthase complexes. In contrast, the architecture of the preprophase band (PPB) and spindle MTs appears more stable, preceding the division plane and direction of chromosomal migration during division (reviewed in Hashimoto, 2015). Interestingly, although MT binding proteins, such as KATANIN (KTN; Luptovciak *et al.*, 2017), SPIRAL2/TORTIFOLIA (SPR2; Buschmann *et al.*, 2004; Shoji *et al.*, 2004; Yao *et al.*, 2008; Nakamura *et al.*, 2018), MICROTUBULE ASSOSCIATED PROTEIN65 (MAP65; Smertenko *et al.*, 2004) have different effects on MT stability and architecture, they do not discriminate between dynamic or more stable MTs and seem to reside on all MT structures found in the cell, including the cortical MT (CMT), the PPB, and the mitotic spindle. Hence it is likely that specific MT binding proteins exist that differentiate between dynamic and stable MT structures in order to allow for these differences in architectural dynamics.

Over the past years, several lines of evidence have shown that specific cues, such as light and auxin are able to influence MT dynamics (Lindeboom *et al.*, 2013; Chen *et al.*, 2014). However, it remains unclear how such cues could be integrated and lead to the required MT orientation. A recent study of AUXIN RESPONSE FACTOR5/ MONOPTEROS (MP) downstream target genes (Möller *et al.*, 2017), identified an overrepresentation of members of the IQ67-domain (IQD) family as being downregulated following impaired auxin response in the early embryo. The founding member of the IQD family, AtIQD1, was shown to bind Calmodulin (CaM), as well as several MT-associated proteins (Levy *et al.*, 2005; Bürstenbinder *et al.*, 2013). A recent systematic analysis of the AtIQD family revealed a diverse array of protein localization and showed that at least some IQD proteins can control MT cytoskeleton topology (Bürstenbinder *et al.*, 2017). Thus, given that both auxin and calcium modulate the MT cytoskeleton (Chen *et al.*, 2014), it is possible that IQD proteins mediate the influences of these signals on the cytoskeleton.

Here, we show that the *IQD15-18* subclade of *Arabidopsis* is evolutionarily conserved throughout the Embryophytes, regulated by auxin and potentially acts as an integrator of different signaling pathways. We further hypothesize that IQD15-18 control MT dynamics through modulating SPR2 activity likely in a CaM dependent way and thereby providing a missing link between these important regulatory pathways.

## Results

### IQ67-domain gene family expression is under developmental and auxin control

Out of the 33 members in the *Arabidopsis IQD* gene family (Abel *et al.*, 2005), a few were reported to be misregulated in embryos in which auxin response was downregulated (Möller *et al.*, 2017). To determine if *IQD* gene regulation is a significant output of auxin action, we carefully examined the expression levels of the entire *IQD* gene family upon reduced auxin signaling. Analysis of two published datasets (one performed on seedlings [Schlereth *et al.*, 2010] and the other on embryos [Möller *et al.*, 2017]) revealed that expression of 13 members (~40% of the family) is misregulated upon inhibition of auxin response (Figure 1A). Several subclades appear to be co-regulated, as is evident from for example the *IQD15-18* clade, that responds similarly in both datasets. To further dissect if *IQD* genes are a direct and rapid auxin output, we assessed the effect of exogenous auxin application on the expression levels of this subclade specifically. Both qPCR and analysis of *promoter::n3*GFP fluorescence intensity after auxin treatment revealed a fast upregulation of *IQD15* transcripts (Figure 1B-C), indicating that this gene is likely a direct target of auxin signaling. Moreover, it was previously shown that *IQD15* expression was reduced in a *mp* mutant background (Möller *et al.*, 2017). Diverse transcriptional response after auxin treatment was evident from qPCR analysis on the other three IQD genes (Figure S1A). Indeed, putative ARF binding sites (AuxREs; Boer *et al.*, 2014) could be identified in close proximity of start codons of all four *IQD* genes (Figure S1B). Taken together, this confirms that auxin is able to regulate the expression of *IQD15* and its close homologs.

**Figure 1:**
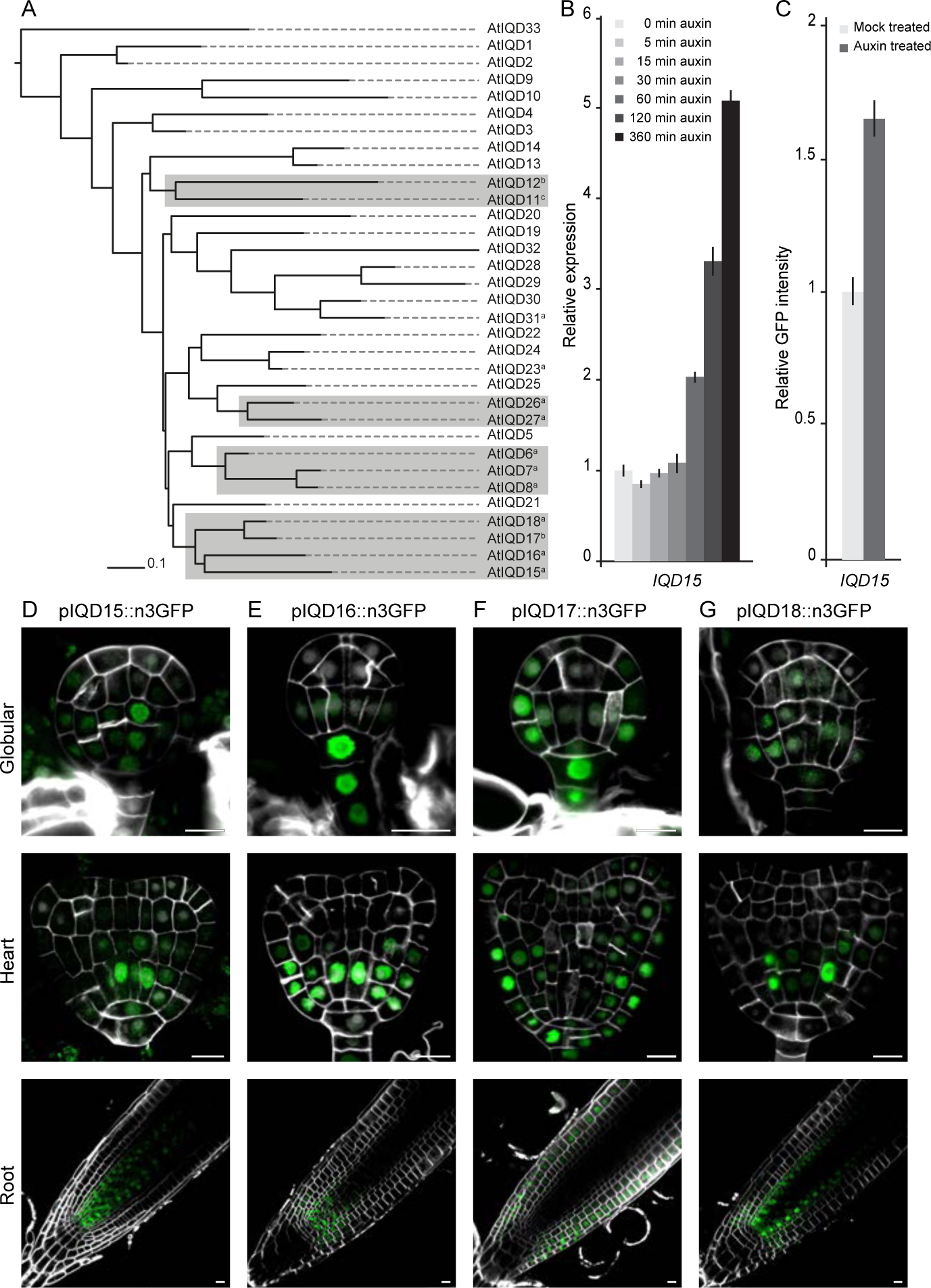
IQD gene expression is under control of auxin signaling. (A) Phylogenetic tree of Arabidopsis IQD proteins, rooted to AtIQD33, shows co-regulation of auxin signaling in several subclades. a= >1.5 fold down in embryo array (Möller *et al.*, 2017); b= >1.5 fold down in seedling array (Schlereth *et al.*, 2010); c= >1.2 fold up in seedling array (Schlereth *et al.*, 2010). (B) Bar diagram showing relative expression of IQD15 in Arabidopsis roots after exogenous auxin treatment for indicated time. (C) Bar diagram showing relative GFP intensity of pIQD15::n3GFP expressing root tips after overnight auxin treatment (1 μM 2-4D). Error bars indicate SEM, n = 3 (B) / 10 (C). (D-G) Expression pattern of IQD15-18 in globular and heart stage embryos and root tips, reveals expression in developmental regions of high auxin signaling. Measuring bar = 10 μM.

The expression profile of the *IQD15-18* subclade was determined by analysis of *promoter::n3*GFP fusion lines throughout *Arabidopsis* developing embryos and primary root meristems. Expression of *IQD15* was observed from globular stage of embryogenesis and stayed restricted to the vascular precursor cells (Figure 1D). In the postembryonic root, expression was also mostly restricted to the vascular tissue and appeared strongest close to the QC (Figure 1D). A similar expression pattern was observed for *IQD18* (Figure 1G), although its expression was somewhat broader in the postembryonic root and expanded into the ground tissue layers and the lateral root cap (Figure 1G). *IQD16* showed a more dynamic expression pattern as it was expressed from four-cell stage of embryogenesis in the suspensor and later (from early heart stage onward) in the entire basal pro-embryo (Figure 1E). In the root, expression could be observed in all tissue types except for QC and columella (Figure 1E), again expression appeared strongest closer to the QC. Finally, *IQD17* was found to complement the expression of *IQD15*, and was expressed in the suspensor of the early globular embryo and the outer tissues of the epidermis and ground tissue (Figure 1F). While similar expression was observed in the root, *IQD17* was excluded from the columella and QC (Figure 1F). Expression could also be observed outside the context of the embryonic and primary root (e.g. lateral roots and developing leaf primordia; Figure S2). Interestingly, expression of *IQD15, −16* and *-18* seems to coincide with regions of high auxin signaling and the observed auxin gradient (Liao *et al.*, 2015), consistent with their regulation by the auxin signaling-pathway.

### IQD proteins mark dynamic microtubule structures

We next generated translational fusion lines of the IQD15-18 subclade to investigate their subcellular protein localization patterns. Proteins could be detected only within domains of promoter activity in both embryos and roots, suggesting that proteins do not move outside their expression domain (Figure 2A-C). Similar to previous reports (Bürstenbinder *et al.*, 2013, 2017), we observed proteins in strands resembling MT-structures and found these structures to be sensitive to treatment with the MT-destabilizing drug oryzalin (Figures 2A-C). In addition to being associated to CMT, IQD18 was also found to reside in the nucleus. Not all cells showed nuclear localized IQD18 and this was especially clear in the postembryonic root where this seemed to be correlated to the developmental age of the cell (Figure 2D), indicating differential localization of IQD18 throughout the cell cycle. Further investigation into possible cell cycle regulation on the localization of IQD18 protein was performed by treatment with hydroxyurea (HU), that stalls the cell cycle in S-phase (Cools *et al.*, 2010). The number of cells with nuclear-localized IQD18 protein was significantly increased after 15 hours of treatment (Figure 3A-C), suggesting re-localization occurs before or during S-phase. Live time-lapse imaging on roots revealed a reduction of IQD18-YFP signal at the lateral sides of the cell to coincide with the appearance of nuclear-localized protein (Figure 3D, white arrowheads; Movie 1). Given that the disappearance of IQD18 protein from the lateral sides and reappearance in the nucleus occurs within minutes (Figure 3D), it is very likely that active protein re-localization takes place. At later stages of the cell cycle, as the nuclear envelope dissolves in (pro)metaphase, IQD18 protein dissipates throughout the cytoplasm, later localizes to the newly forming cell plate, and can occasionally be observed in spots in the daughter nuclei (Figure 3E-F and Movies 2 and 3). The dynamic localization of IQD18 was observed in multiple cell types, indicating a cell type-independent regulation on localization of IQD18 protein through the cell cycle.

**Figure 2:**
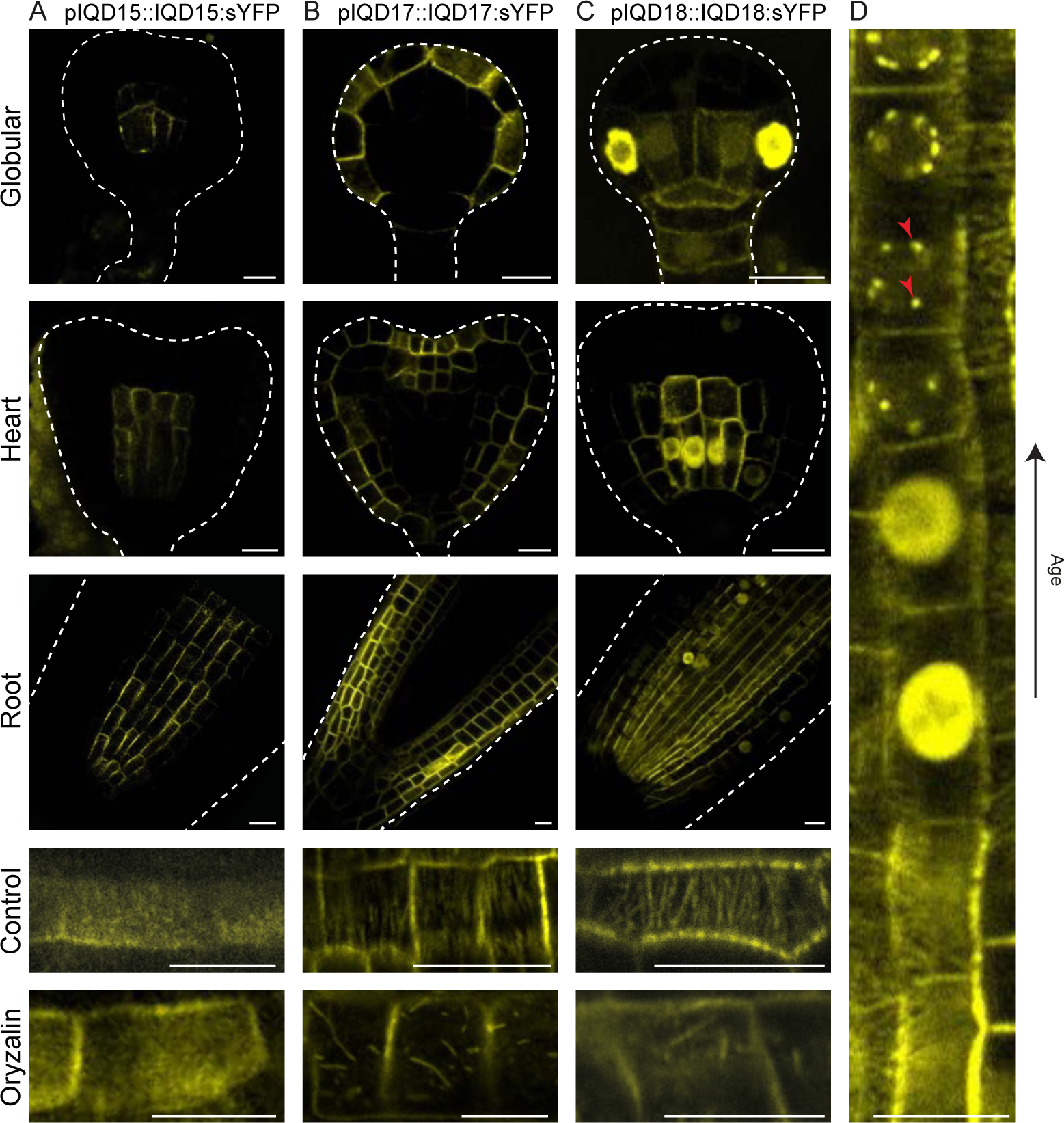
IQD proteins localize to microtubules in vivo. Subcellular protein localization of IQD15 (A), −17 (B), and −18 (C), in globular and heart stage embryos and root tips reveals oryzalin sensitive microtubule association within expression domain. White dashed line represents embryo and root outline. (D) Dynamic protein localization of IQD18 in root tips suggests cell cycle dependent subcellular localization. Measuring bar = 10 μM.

**Figure 3:**
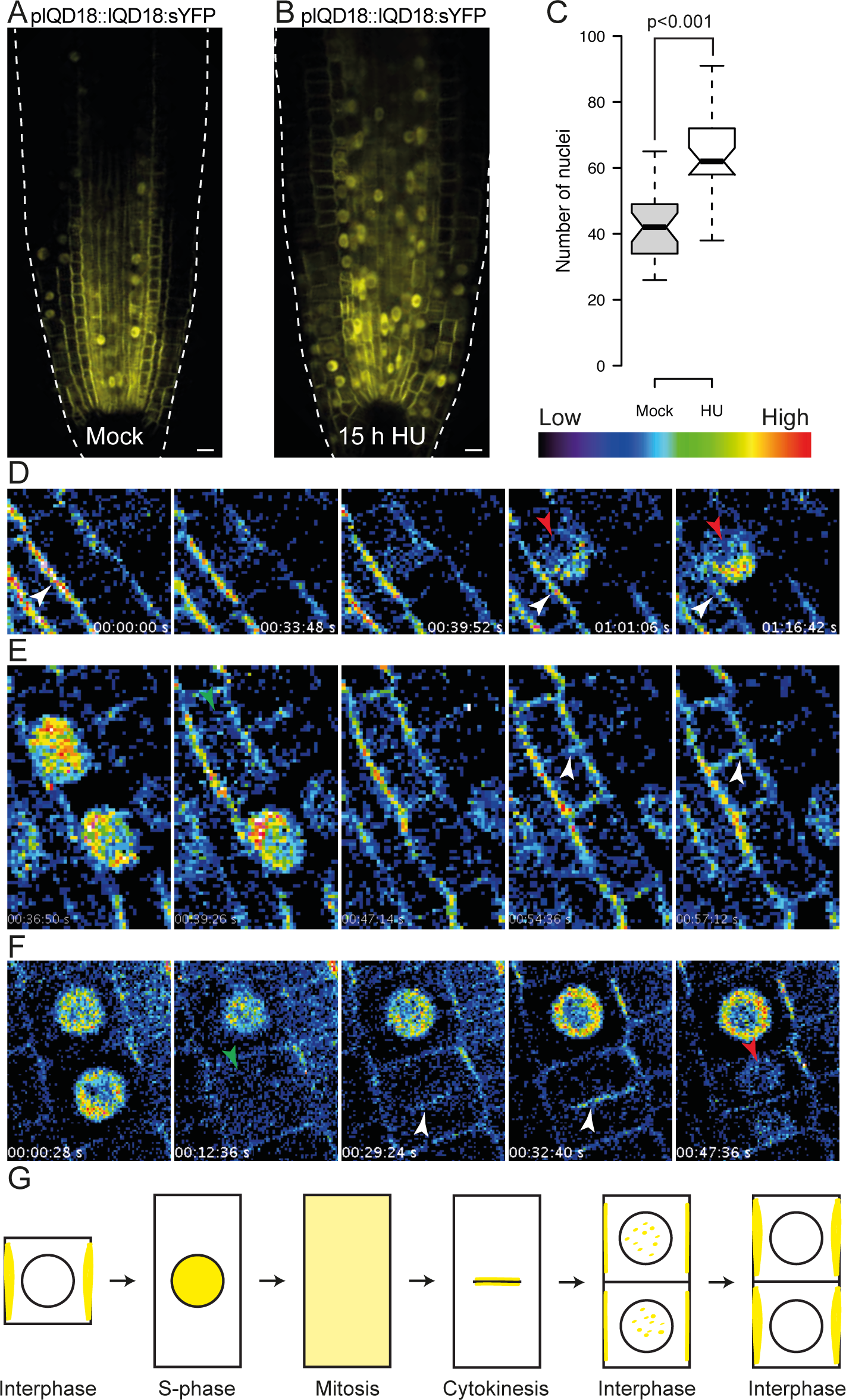
Dynamic IQD18 protein localization is cell cycle dependent. (A-C) Cell cycle arrest in S-phase (B) results in significantly increased cells with nuclear localized IQD18 protein, compared to control conditions (A). n = 29; measuring bar = 10 μM. (D-F) Time-lapse imaging of IQD18 protein becoming nuclear in S-phase (D) and in endodermal (E) and cortical cells (F) through division, reveals dynamic subcellular localization. (G) Schematic representation of dynamic IQD18 protein localization through different phases of cell cycle.

As protein localization between the different members of this same subclade differs, with only IQD18 displaying nuclear accumulation, we were interested to know the ancestral localization mode within this subclade. We investigated the occurrence of *Arabidopsis thaliana* IQD15-18 (AtIQD) orthologues in a number of different species, including tomato, poplar, rice, maize, and moss (Figure 4A). All analyzed monocot species possessed only a single copy of this subclade, while all eudicots (with the exception of Medicago) had multiple, suggesting a multiplication event of this subclade occurred in eudicots (Figure 4A). This would also suggest the localization properties of a monocot orthologue to be similar to that of a common ancestor. To test this, we generated a GFP protein fusion of the rice orthologue *OsIQD14*, which has up to 61% similarity to the *Arabidopsis* genes and shares highly conserved domains (Figure S3). Interestingly, in both rice and *Arabidopsis* roots, this protein localized in a pattern that was indistinguishable from that of AtIQD18, with clear microtubule association, nuclear localization, and similar behavior during the cell cycle (Figure 4B-D). This strongly suggests that AtIQD18 has retained its ancestral localization properties while other members of the subclade have diversified.

**Figure 4:**
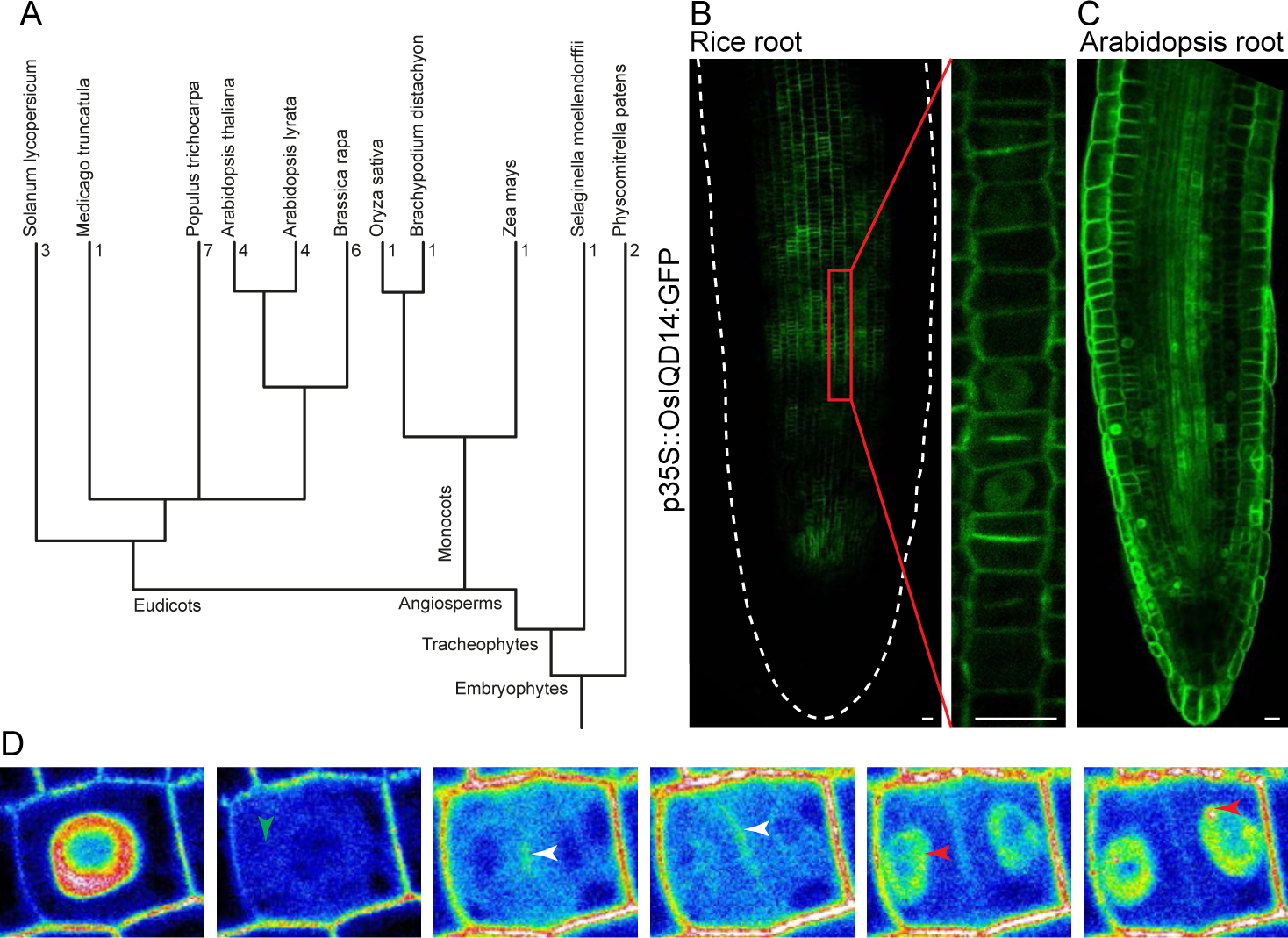
IQD sequence and protein localization is evolutionarily conserved. (A) Phylogenetic tree of land plants annotated with number of AtIQD15-18 co-orthologs per species, shows multiplication occurring after monocot/dicot split. (B-D) Subcellular protein localization of p35s::OsIQD14:GFP in rice (B) and Arabidopsis (C) root tips and through cell division in Arabidopsis (D) reveals identical localization as AtIQD18. Measuring bar = 10 μM.

Generally, MT-associated proteins mark both the highly dynamic cortical MT network and seemingly more stable MT structures like the PPB and the mitotic spindle (e.g. Marc *et al.*, 1998; Bao *et al.*, 2001; Buschmann *et al.*, 2004; Smertenko *et al.*, 2004). Interestingly, none of the IQD proteins in the IQD15-18 clade in *Arabidopsis*, nor the rice OsIQD14 protein associated with either the PPB or the mitotic spindle (Figures 2, 3, 4 and S4 and Movies 1-3). Thus, these proteins preferentially associate with dynamic MT structures.

### IQDs directly bind MT’s and interacts with Calmodulin and SPIRAL2 in vivo

Association of proteins to MT can either be through direct protein-protein interaction or could alternatively be bridged by MT-binding proteins. To determine if IQD15-18 represent Microtubule-Associated Proteins (MAPs), we tested direct MT binding capabilities of AtIQD18 using an *in vitro* MT Binding Protein Spin-Down assay using recombinant AtIQD18 and purified MT. This showed a clear co-sedimentation of recombinant AtIQD18 with MT (Figure 5A), similar to the positive control (MAP2). Together with similar results we obtained from OsIQD14 (Yang *et al.*, 2018), this indicates an evolutionarily conserved property of IQD proteins to directly bind MT.

**Figure 5:**
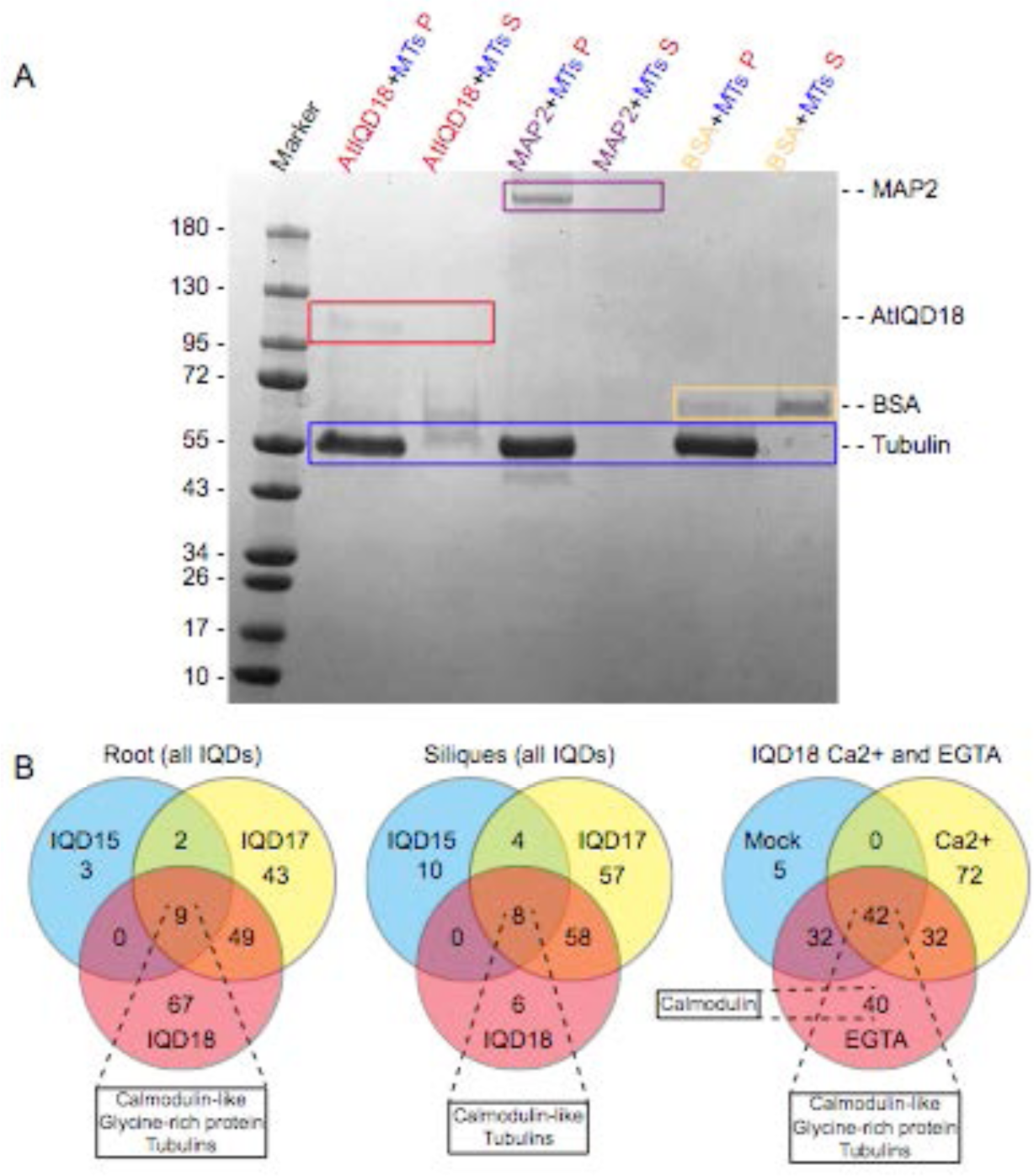
IQD protiens directly bind microtubules in vitro and associate with Calmodulins in vivo. (A)in vitro MT binding assay reveals direct binding between AtlQD18 and,similar to positive control MAP2.AtlQD18 is outlined in red, positive control MAP2 in purple negative control BSA in yellow and Tubulin in blue.P=pellet fraction; S=supernatant fraction.(B) Venn diagrams portraying IP-MS/MS experiments on AtlQD15, −17,and −18 roots and siliquess, reveal overlapping interactions with Calmodulins and tubulins.

We next used an *in vivo* approach to identify interaction partners of AtIQDs in different developmental contexts. We performed several immunoprecipitation experiments followed by tandem-mass spectrometry (IP-MS/MS) on siliques (embryo context) or roots of plants expressing AtIQD15, −17, and −18 fusion proteins under control of their endogenous promoter. As previously reported for IQD1 (Bürstenbinder *et al.*, 2013), we found Calmodulin (CaM) and Calmodulin-like (CML) proteins as well as Tubulin to associate with all three IQD proteins tested (Figures 5B and S7, Table 1 and S1). We next used Yeast-Two-Hybrid (Y2H) and Bimolecular Fluorescence Complementation (BiFC) to determine if interactions with these CaM/CML proteins are based on direct protein-protein interactions. Y2H showed that both IQD17 and IQD18 directly interact with different CaM proteins (Figure S5). BiFC confirmed the interaction between IQD18 and CaM1 (Figure S6), and furthermore showed that this interaction occurs at MT structures (Figure S6A’ and B’). Since CaM proteins are not by themselves known to interact with MTs (Bürstenbinder *et al.*, 2013), it is likely that IQD18 recruits CaM1 to MTs, a property that we also observed for OsIQD14 (Yang *et al.*, 2018).

**Table 1:**
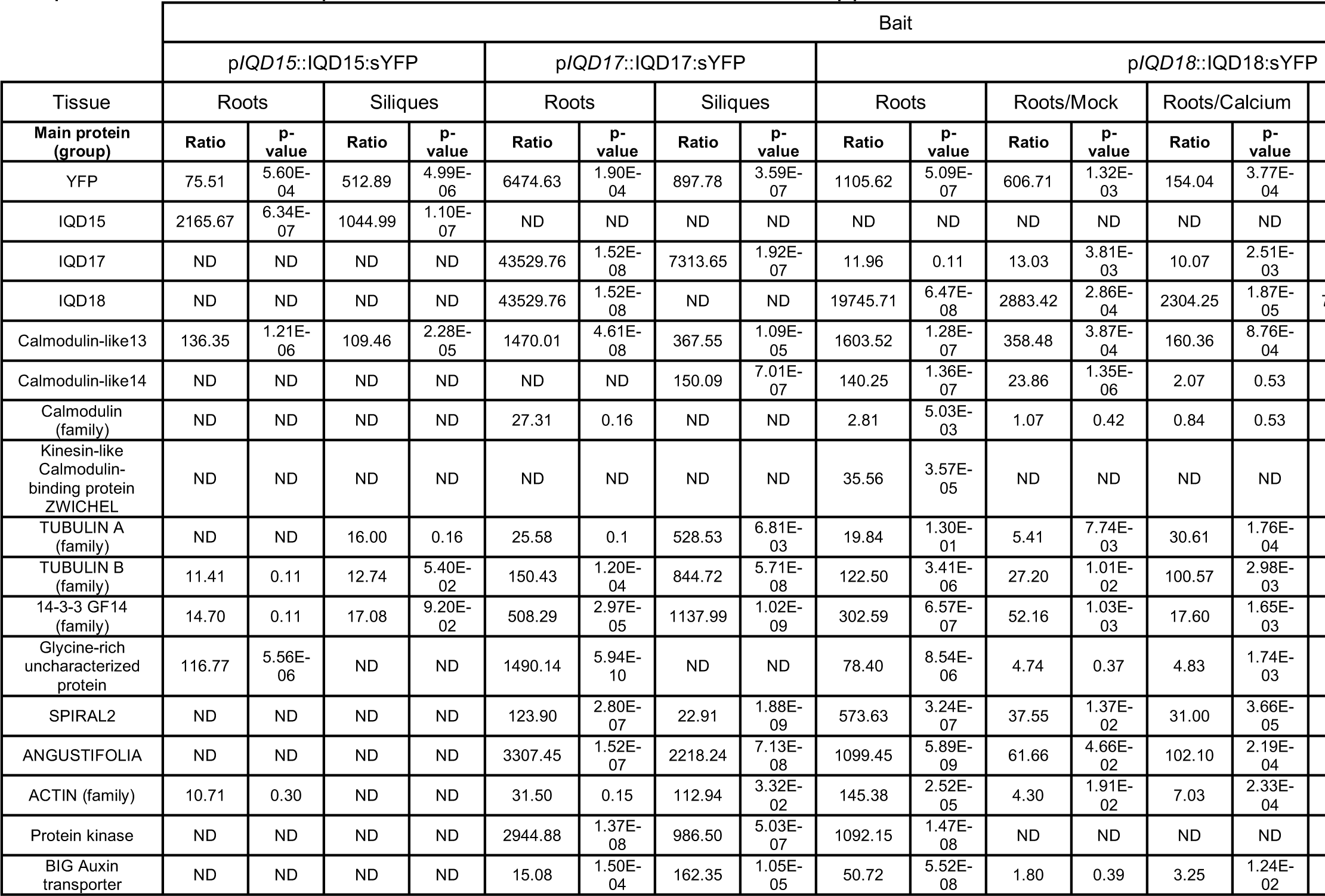
Overview of selected protein groups identified through IP-MS/MS. Selected identified main protein groups are shown, ratios were calculated compared to the wild-type control and p-values by means of a Student’s t-test (n=3); Stickiness is calculated from the occurrence of the identified protein group among an unpublished dataset of 86 independent IP-MS/MS experiments; ND = Not Detected; NA = Not applicable

In addition to confirming MT and CaM/CML interactions, our IP-MS/MS experiments identified a range of novel IQD-interacting proteins. These included a Glycine-rich protein, a kinesin, ANGUSTIFOLIA, and several members of the 14-3-3 type GF14 proteins (Figures 5B and S7, Table 1 and S1). Interestingly, IQDs were also found to bind the SPIRAL2 (SPR2) protein (Table 1). SPR2 was recently found to bind the minus-end of MT and is characterized by its loss-of-function phenotype of spiraling tissues (Buschmann *et al.*, 2004; Shoji *et al.*, 2004; Wightman *et al.*, 2013; Nakamura *et al.*, 2018; Leong *et al.*, 2018). IQD18 and SPR2 proteins directly interacted in Y2H (Figure S8A) and BiFC assays showed that also these interactions occurred at MT structures (Figure S8B).

SPR2 is an important regulator of MT dynamics, but no interactors or regulators have so far been identified. To determine if the interactions revealed in IQD IP-MS/MS are representative of SPR2 protein function, we also carried out similar experiments with p*35S*::SPR2:GFP (Shoji *et al.*, 2004). This revealed an overlapping interactome between IQDs and SPR2, including several of the same CaMs/CMLs and 14-3-3 GF14 proteins (Table S1 and Figure S9). We did not find any IQD proteins, but might be due to their low protein abundance and tissue specific expression, compared to ubiquitous SPR2 protein accumulation.

### Calcium modulates the assembly of MT complexes

Given that IQD proteins interact directly with several CaM/CML proteins, and that this interaction likely recruits CaM/CML proteins to MTs, we asked if calcium would affect the IQD protein interactome. We therefore performed independent IP-MS/MS experiments on roots of a IQD18-YFP line in high or low calcium conditions (by addition of either 100 mM CaCl_2_ or 20 mM EGTA, respectively). Through statistical analysis, we identified several proteins that show differential binding in either condition (Table 1 and S1 and Figure S10). Among these proteins, we also found several CaM proteins that showed more prominent binding in low-calcium conditions (Table 1). This suggests that calcium, presumably through binding to CaM/CML proteins, modulates their ability to bind IQD proteins. We tested this directly by quantifying the binding of recombinant IQD18 protein to CaM-containing beads in the presence of buffer, EGTA or calcium. This showed that indeed, calcium increased the binding of IQD18 to this generic model CaM protein, showing that IQD-CaM interactions can be modulated by calcium in a reconstituted *in vitro* system (Figure S11). Hence, the calcium-induced changes in CaM/CML protein association in the IP-MS/MS experiments are likely also mediated by direct effects of calcium on CaM/CML proteins. Importantly, not only the IQD-CaM/CML interactions were affected by calcium and EGTA treatment, also the association of SPR2 was modulated in these experiments (Table 1; Yang *et al.*, 2018). This reveals new cellular effects of calcium in plant cells: recruitment of CaM/CML to MTs, and modulation of the IQD-SPR2 association.

### Nuclear localization of IQD proteins is important for proper calcium responses

Among the IQD15-18 clade, IQD18 is the most dynamic in its subcellular localization, a property that is shared with its rice orthologue OsIQD14 (Figures 3 and 4). It is likely that this regulated localization is important for protein function, and we used a misexpression strategy to alter IQD18 protein levels and localization. We first generated a line expressing this protein under control of the strong meristematic *RPS5A* promoter (Weijers *et al.*, 2001) and fused with sYFP (p*RPS5A*::IQD18:sYFP; R18). This line showed similar localization patterns as observed in the genomic fusion line, with CMT and nuclear localized protein, in both embryos and root tips (Figure 6A). We did not observe any obvious phenotypic changes in these misexpression lines in embryos nor in roots. We then cultured seedlings in EGTA to limit the endogenous calcium levels. This treatment strongly inhibited root growth in wild-type plants, and we found that R18 lines were more sensitive to EGTA-induced root growth inhibition (Figure 6E).

**Figure 6:**
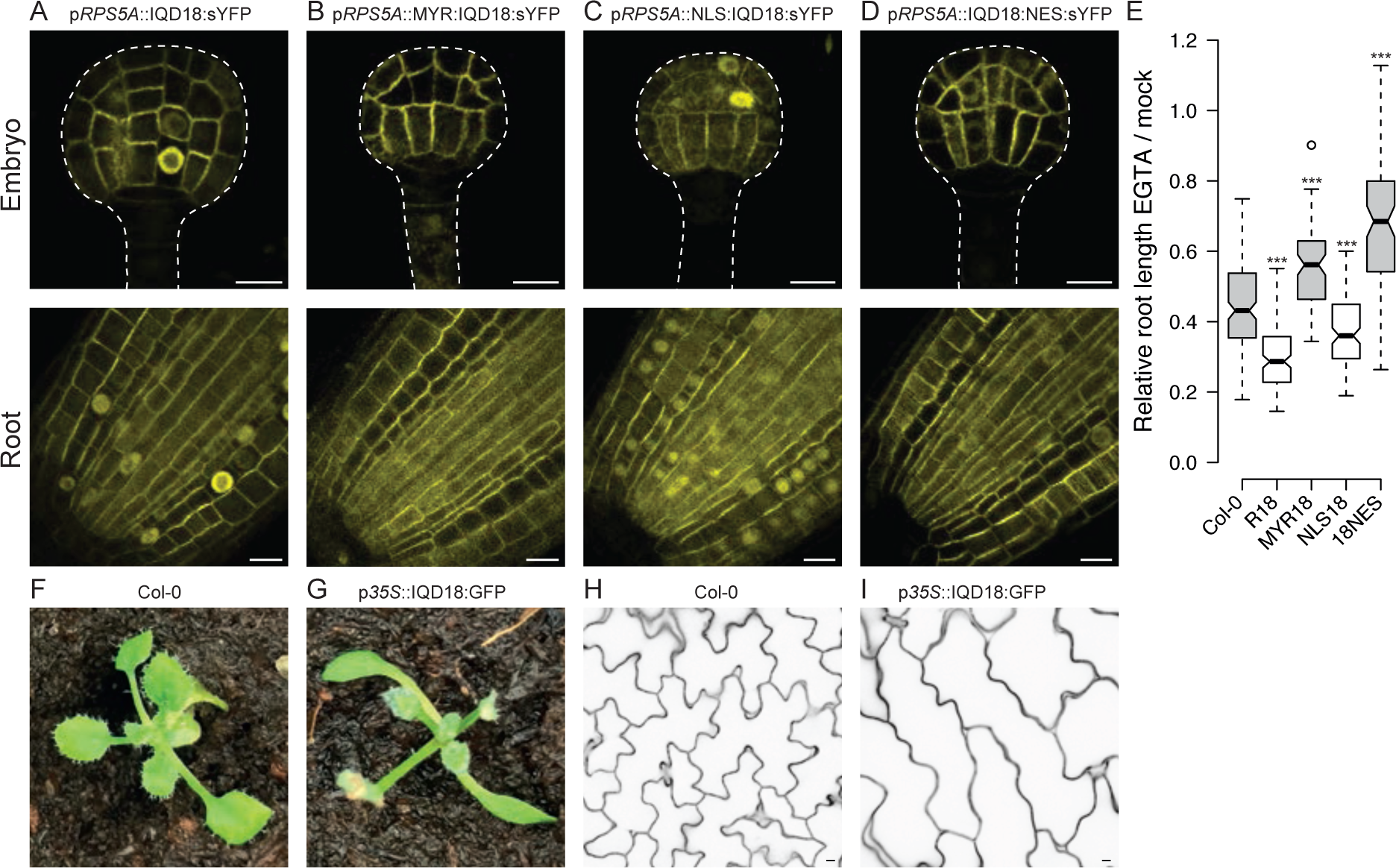
Nuclear localization of AtIQD18 is important for proper calcium signaling and misexpression leads to spr2-like phenotypes. (A-E) Misexpression and mislocalization of AtIQD18 by means of RPS5A promoter (A) and myristoylation (B; MYR), nuclear localization (C; NLS), and nuclear export (D; NES) tags, results in altered calcium signaling and reveals a role for nuclear localization in this process (E). (F-J) Overexpression of AtIQD18 results in spr2-like phenotypes of spiraling cotyledons (G) compared to wild-type (F) and stretched and less complex cotyledon pavements cells (I-J). Measuring bar = 10 μM.

We next exploited the increased sensitivity to EGTA as a measure for biological activity of IQD18 protein, and asked which subcellular localization is associated with activity. We generated *Arabidopsis* lines expressing fusion proteins with three different localization tags to alter the subcellular localization of the AtIQD18 protein. We made use of an N-terminal fused myristoylation tag (MYR; Traverso *et al.*, 2013) for membrane anchoring (p*RPS5A*::MYR:IQD18:sYFP; MYR18), an N-terminal nuclear localization signal (NLS; Lange *et al.*, 2007) for increased nuclear targeting (p*RPS5A*::NLS:IQD18:sYFP; NLS18), and a C-terminal nuclear export signal (NES; Gallagher and Benfey, 2009) for export out of the nucleus (p*RPS5A*::IQD18:NES:sYFP; 18NES). All tags appeared to function properly, as we observed increased subcellular localizations at the expected sites in all lines. Interestingly, while neither the NLS or the NES conferred exclusive localization in- or outside the nucleus, membrane anchoring by means of the MYR-tag seemed to fully localize AtIQD18 to the membrane and completely abolished nuclear localization (Figure 6B-D).

Both lines with increased nuclear AtIQD18 (i.e. R18 and NLS18) showed EGTA hypersensitivity, while lines with no or reduced nuclear-localized AtIQD18 (i.e. MYR18 and 18NES), showed EGTA resistance (Figure 6E). This suggests that nuclear localization of this protein is important for proper calcium responses in the root. IQD15-18 genes are expressed most prominently in young, dividing tissues, which overlaps mostly with the activity domain of the *RPS5A* promoter (Weijers *et al.*, 2001). To misexpress IQD18 outside of its normal expression domain, we next expressed AtIQD18 protein fused to GFP, under control of the Cauliflower mosaic virus *35S* promoter. Similar to what was previously reported for AtIQD16 (Bürstenbinder *et al.*, 2017), we observed strong phenotypes in these lines (Figure 6F). Specifically, cotyledons showed strong spiraling (Figure 6G). Interestingly, this phenotype strongly resembles the spiraling phenotype observed in mutants that affect the MT cytoskeleton (Hashimoto, 2002), including SPR2 (Furutani *et al.*, 2000; Buschmann *et al.*, 2004; Shoji *et al.*, 2004). This suggests that IQD proteins control the MT cytoskeleton, presumably through their interaction with SPR2.

Together, these results strongly indicate that IQD proteins are a new class of proteins that can directly bind and affect MT. With a wide array of interacting proteins, they could function as integrator of signals, regulating MT dynamics.

## Discussion

Auxin has a profound effect of the organization of the CMT is plant cells, however, how auxin mechanistically effects and generates potential for dynamic MT organization, remains unknown. Here we provide evidence for a missing link that can connect auxin signaling to downstream MT modulators, through an evolutionarily conserved subclade of the IQD family as integrator of different signals. While we were unable to find single loss of function mutants that showed altered development, likely due to redundant action of the studied subclade (data not shown), mutations in the single rice orthologue have significant effects on grain size and quality (Yang *et al.*, 2018), showing the importance of this protein family for development. Recent studies on the family of IQ67-domain proteins (Bürstenbinder *et al.*, 2017; Sugiyama *et al.*, 2017) have shed light on their potential to alter MT stability and organization in different contexts. We show that the subclade of *AtIQD15-18* genes is transcriptionally regulated by auxin signaling and that these genes are expressed in the developmental regions that correspond to high signaling. Furthermore, we show that the IQD proteins can directly bind to CaM and MT *in vitro* and that they preferentially localize to the dynamic CMT structures, *in vivo*. The interaction between IQD and CaM was found to be modulated by calcium. Interestingly, co-overexpression of a CaM with OsIQD14 was recently found to suppress IQD function and restore its phenotype (Yang *et al.*, 2018). This suggests that calcium signaling is able to control the interactions of IQD proteins and thereby modulate their function. Calcium signaling was in turn found to be dependent on correct localization of IQD proteins, as altering the nuclear localization properties of IQD18 resulted in impaired responses. Although the precise role of nuclear-localized IQD protein remains unclear, this could involve parts of the nuclear calcium signaling (Charpentier and Oldroyd, 2013), as for example interactions with a nuclear ion channel were also observed (Table S1). Moreover, we identified novel IQD- interacting proteins, including the MT minus-end binding protein SPR2. We were able to confirm that this is a direct interaction and also that this interaction can be modulated by calcium, which suggests an auxin-calcium-IQD-SPR2 pathway could be controlling MT dynamics. In our efforts to confirm IQD-SPR2 interaction we identified novel interactors of SPR2, including a phosphatase and a kinase, in addition to an overlapping interactome between IQD and SPR2 (Table S1). Considering that phosphorylation was proposed as a regulatory mechanism for SPR2 (Wightman *et al.*, 2013), this provides valuable insights for future research directions on the regulation of this protein.

We propose a model for how auxin mediates MT dynamics and organization, through the action of IQD proteins (Figure 7). Through (slow) transcriptional regulation on the expression of *IQD* genes, auxin potentiates control on the MT organization. Calcium levels are well known to increase after an auxin peak (Monshausen *et al.*, 2011; Monshausen, 2012) and this will mediate fast auxin-dependent regulation on MT organization: In resting conditions (low calcium concentrations), IQD proteins bind to SPR2 and thereby inhibiting its function to stabilize MT minus ends (Nakamura *et al.*, 2018; Leong *et al.*, 2018), leading to higher stability of the MT structure (less dynamicity). In an event of an auxin peak, calcium levels will rise leading to higher affinity between CaM and IQD (Yang *et al.*, 2018 and this study), which will free SPR2 to bind MT minus ends and thereby increasing stability of branching MT and increasing dynamicity of the MT architecture. This model is further supported by the finding that overexpression of CaM represses IQD function, as shown by a rescue of the *spr2*-like phenotype (Yang *et al.*, 2018). Under normal physiological conditions, these processes will be highly coordinated and slight differences in concentration of signaling molecules will locally affect binding affinities between the different components and thus have a fine-tuned effect on MT organization. In this way, IQD proteins contribute to the integration of signals from both auxin and calcium signaling to modulate the MT dynamics and architecture, affecting for example rice grain size and quality (Yang *et al.*, 2018). What remains unclear and will be interesting for future studies, is how the cell cycle-dependent subcellular localization of these proteins contributes to this process, what function these proteins play during *Arabidopsis* development, as well as how the diverse localization of the Arabidopsis IQD15-18 proteins are integrated in a redundant function.

**Figure 7:**
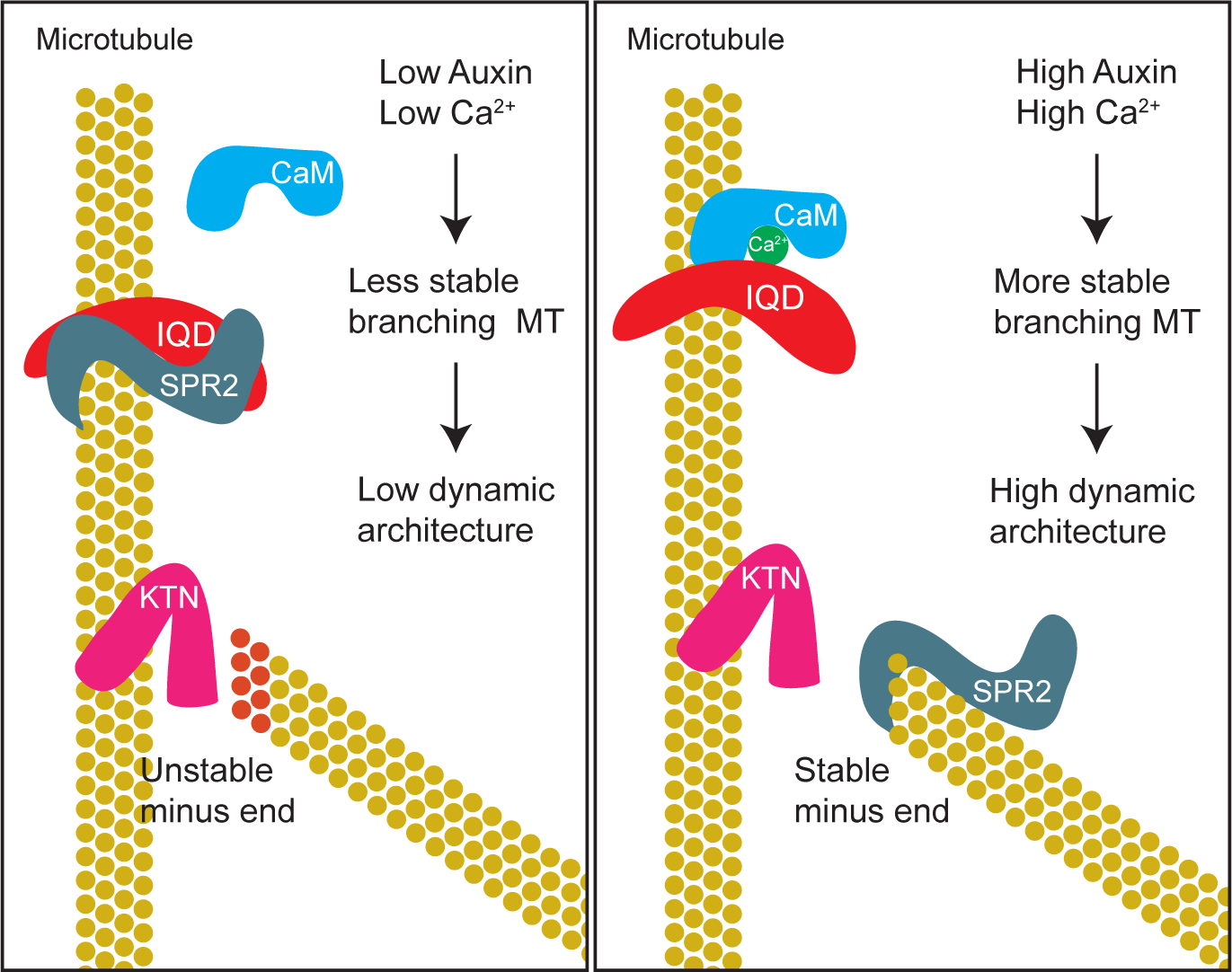
Proposed model for IQD function on MT dynamics.

## Methods

### Genome mining and phylogenetic tree assembly

Multiple sequence alignment was performed on protein sequences of all (full length) *Arabidopsis* IQD proteins, and a phylogenetic tree was assembled using only non- gap generating sequences, using MAFFT (Katoh and Standley, 2013) and AtIQD33 was used to root the tree. Protein sequences of AtIQD15-18 were used as query in a BLAST to find related proteins in transcriptome databases of different species (i.e. *Solanum lycopersicum, Medicago truncatula, Populus trichocarpa, Arabidopsis lyrata, Brassica rapa, Oryza sativa, Brachipodium distachyon, Zea mays, Selaginella moellendorfii*, and *Physcomitrella patens*). Reciprocal BLAST on *Arabidopsis* protein database was used to filter the recovered hits and only those hits that resulted in AtIQD15-18 as top hits were kept.

### Plant materials and growth conditions

Previously described plant lines expressing p*35S*::SPIRAL2:GFP (Shoji *et al.*, 2004) and p*35S*::OsIQD14:GFP (Yang *et al.*, 2018) were used. Seeds were surface sterilized and grown on ½ MS (Duchefa) plates under standard continuous light growth conditions at 21°C following a one to four-day stratification at 4°C. *Arabidopsis* ecotype Columbia-0 was used as wild-type control in all cases. Chemical and hormone treatments were performed by either germinating seeds on supplemented media or transferring seedlings from normal media to supplemented media and continuing growth for indicated time.

### Cloning and plant transformation

Promoter fragments (up to 5 kb upstream of start codon), translational genomic fusions (promoter fragment plus coding genomic fragment) and coding sequences (CDS) were amplified from genomic DNA (promoter and genomic) or root cDNA (CDS) using PCR and Phusion Flash master mix (Thermo Scientific) or Q5 DNA polymerase (New England BioLabs) and the primers described in Table S2. PCR products were cloned into the pPLV4_v2 (promoter) or pPLV16 (genomic) vectors using Ligation Independent Cloning (LIC; De Rybel *et al.*, 2011; Wendrich *et al.*, 2015a). For misexpression and mislocalization coding sequences were cloned into pPLV28 using LIC (De Rybel *et al.*, 2011; Wendrich *et al.*, 2015a) or pDONR221 using Gateway cloning (Karimi *et al.*, 2007) and different mislocalization tags were added during PCR using primers listed in Table S2. All constructs were confirmed by Sanger sequencing and transformed into *Arabidopsis* Col-0 wild-type plants through Agrobacterium mediated transformation. At least three independent transformants were checked and representative pictures are shown.

### Microscopic analysis

Confocal Laser Scanning Microscopy (CLSM) was performed, as described previously (Llavata *et al.*, 2013; Wendrich *et al.*, 2015b), on Leica SP5 and SP8X (CLSM) systems. Five-day-old seedling roots were stained with propidium iodide (PI) for 2 minutes and imaged by excitation at 488nm or 514nm and detection 600-700nm (PI). Embryos were examined by isolating ovules and fixing them in a 4% paraformaldehyde / 5% glycerol in PBS solution containing 1.5% SCRI Renaissance Stain 2200 (R2200; Renaissance Chemicals, UK), before extruding embryos and imaging R2200 at 405nm excitation and detection between 430- 470nm. GFP, YFP and RFP were respectively imaged by excitation at 488nm, 515nm or 561nm and detection at 500-535nm, 535-600nm or 600-700nm.

### RNA extraction and qRT-PCR

RNA was extracted from five-day-old *Arabidopsis* seedlings or seedling roots using TriZol (Invitrogen) and subsequently subjected to column purification using an RNeasy Plant kit (Qiagen) following manufacturer’s instructions. Normalized amounts of RNA were used to synthesize cDNA using an iScript kit (Bio-rad). Relative expression of target genes was measured by quantitative-real-time-PCR (qPCR), the primers listed in Table S2, and expression values were normalized against *ACTIN2* and *EEF1*, using qBase software (Hellemans *et al.*, 2007).

### Yeast-Two-Hybrid (Y2H)

Interaction of AtIQD17, AtIQD18 and SPR2 (or CaM1/2/3) was detected by standard yeast two-hybrid analysis following the manufacturer’s instructions (Clontech). cDNAs encoding IQD18, SPR2, CaM1 were subcloned into pGBKT7 and pGADT7 vector, resulting in the fusion of IQD18-AD, SPR2-AD, CaM1-AD, SPR2-BD and CaM1-BD respectively (AD, activating domain; BD, binding domain). Yeast transformants were spotted on the restricted SD medium (SD-Leu/-Trp, short as SD-L/T) and selective medium (SD-Leu/-Trp/-His/, short as SD-L/T/H).

### Bimolecular fluorescence complementation (BiFC)

For BiFC (Bimolecular Fluorescence Complementation) assay, cDNAs encoding *IQD18, SPR2* or *CaM1* were cloned into p*35S*:YFPN or p*35S*:YFPC vector by gateway LR reaction, resulting in constructs expressing IQD18-cYFP, SPR2-nYFP, CaM1-nYFP, respectively. Resultant constructs with control blank vectors were co-expressed in *N. benthamiana* leaves and yellow fluorescence was observed by Leica SP8 confocal microscope using an argon laser excitation wavelength of 488 nm after infiltration for 3 days.

### Recombinant Expression of AtIQD18 and microtubule spin down assay

Coding sequence of AtIQD18 was amplified by PCR (primers *IQD18-COLD-P1/2*) and subcloned into pCold-HF for expression of IQD18-His fusion protein. After confirmation by sequencing, the construct was transformed into *E. coli* BL21(DE3) cells and expression of the fusion protein was induced by adding isopropyl-β-D-thiogalactoside (IPTG; final concentration 1mM) at 16°C overnight. Cells were lysed by sonication in lysis buffer (50 mM NaH_2_PO_4_, 300 mM NaCl and 10 mM imidazole, pH 8.0) and AtIQD18-His protein was purified using Ni-NTA His Bind Resin (Novagen) according to the manufacturer’s protocols.

*In vitro* microtubule-binding assay was performed using Microtubule Binding Protein Spin-down Assay Kit (Cytoskeleton). Briefly, 5 μg purified AtIQD18-His, MAP2 (positive control) and BSA (negative control) proteins were respectively incubated with 10 μg prepolymerized bovine brain tubulin in general tubulin buffer (80 mM PIPES, pH 7.0, 2 mM MgCl_2_, and 0.5 mM EGTA) containing 20 μM taxol. Following centrifugation at 100,000 x g, both soluble and pellet fractions were analyzed by SDS-PAGE and Coomassie Brilliant Blue staining.

### Calmodulin binding assay

Calmodulin binding assay were performed as described before (Levy *et al.*, 2005) with some modifications.

For expression of CaM4-GST, a full-length cDNA fragment encoding the CaM4 was cloned into pDEST-GST using Gateway (Invitrogen). The recombinant CaM4 protein was expressed in BL21(DE3) at 30 °Cfor 4 h by induction with 1 mM IPTG. Bacterial cells were harvested and sonicated in Lysis buffer (50mM Tris-HCL, 150mM NaCl). After centrifugation, the supernatant was used for incubating with GST agroase.

Aliquots of 100 μL of CaM4-GST beads, pre-equilibrated with Lysis buffer, were mixed with 500 μL of bacterial supernatant supplemented with 2 mM CaCl2 or 5 mM EGTA and incubated for 1 h at 4 °C under gentle shaking. CaM4-GST beads were sedimented by centrifugation and washed four times with 500 μL of Lysis buffer, followed by a final wash with 100 μL of the same solution. The bound proteins were eluted by boiling the beads for 2 min in 100μL of 4x SDS sample buffer. Proteins of the total extract, the initial supernatant, the last wash, and the pellet fraction were analyzed by SDS-PAGE and western blot by His antibody.

### Immunoprecipitation followed by tandem mass-spectrometry (IP-MS/MS)

IP-MS/MS was performed, as previously described by Wendrich *et al.* (2017), on up to 3 grams of siliques and five-day-old seedling roots of transgenic *Arabidopsis* plants harboring translational fusion constructs of p*IQD15*::gIQD15:sYFP, p*IQD17*::gIQD17:sYFP, p*IQD18*::gIQD18:sYFP, and whole seedlings harboring p*35S*::SPR2:GFP (Shoji *et al.*, 2004). The same material from Col-0 wild-type plants was collected as control sample. Each sample was performed in triplicate for follow-up statistical analysis. Calcium and EGTA treatments were performed by respectively adding 100 mM and 20 mM during the protein extraction phase.

### Image processing and measurements

All image measurements were performed using ImageJ software and this software was also used for further processing of images. Brightness and contrast levels were globally adjusted and images were cropped and placed on a matching background for ecstatic reasons.

## Acknowledgements

The authors would like to acknowledge Daniël Van Damme and Steffen Vanneste for stimulating discussions; Jan Willem Borst and the MicroSpectroscopy Centre Wageningen for microscopy equipment and assistance; Sjef Boeren and VIB’s Proteomics Expertise Center (PEC) for MS analysis. Work in the lab of D.W. was supported by European Research Council Starting Grant CELLPATTERN 281573; B.D.R. was funded by The Research Foundation - Flanders (FWO; Odysseus II G0D0515N and 12D1815N) and Netherlands Organization for Scientific Research (NWO; VIDI 864.13.00).

---

**Figure S1:**
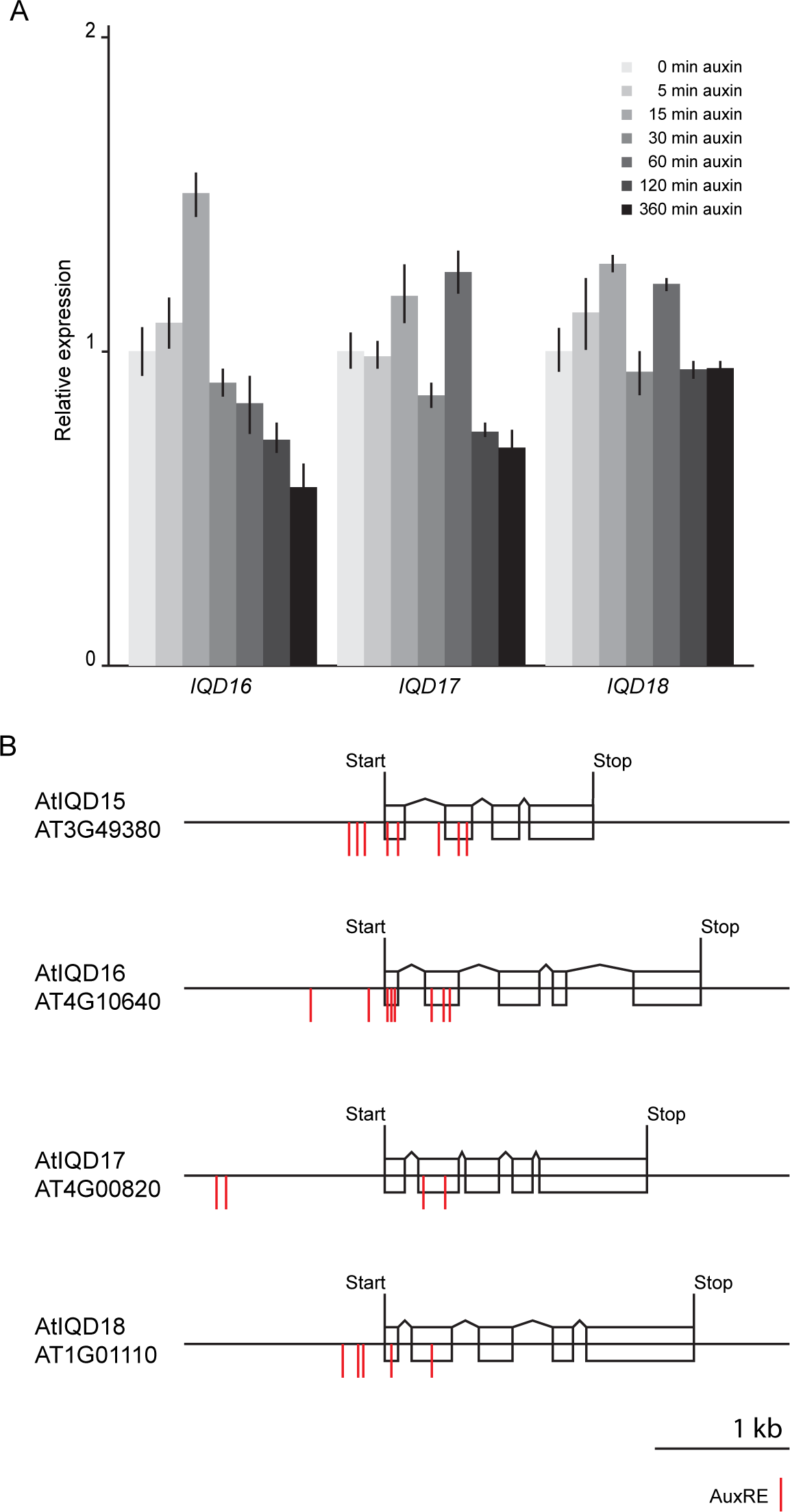
Potential for auxin regulation on IQD genes. (A) Bar diagram showing relative expression of AtIQD16-18 transcripts in Arabidopsis roots after exogenous auxin treatment for indicated time. (B) Schematic representation of genomic regions of AtIQD15-18. Blocks represent exons, lines connection the blocks represent introns. Red lines indicate possible auxin responsive elements close to the translational start site.

**Figure S2:**
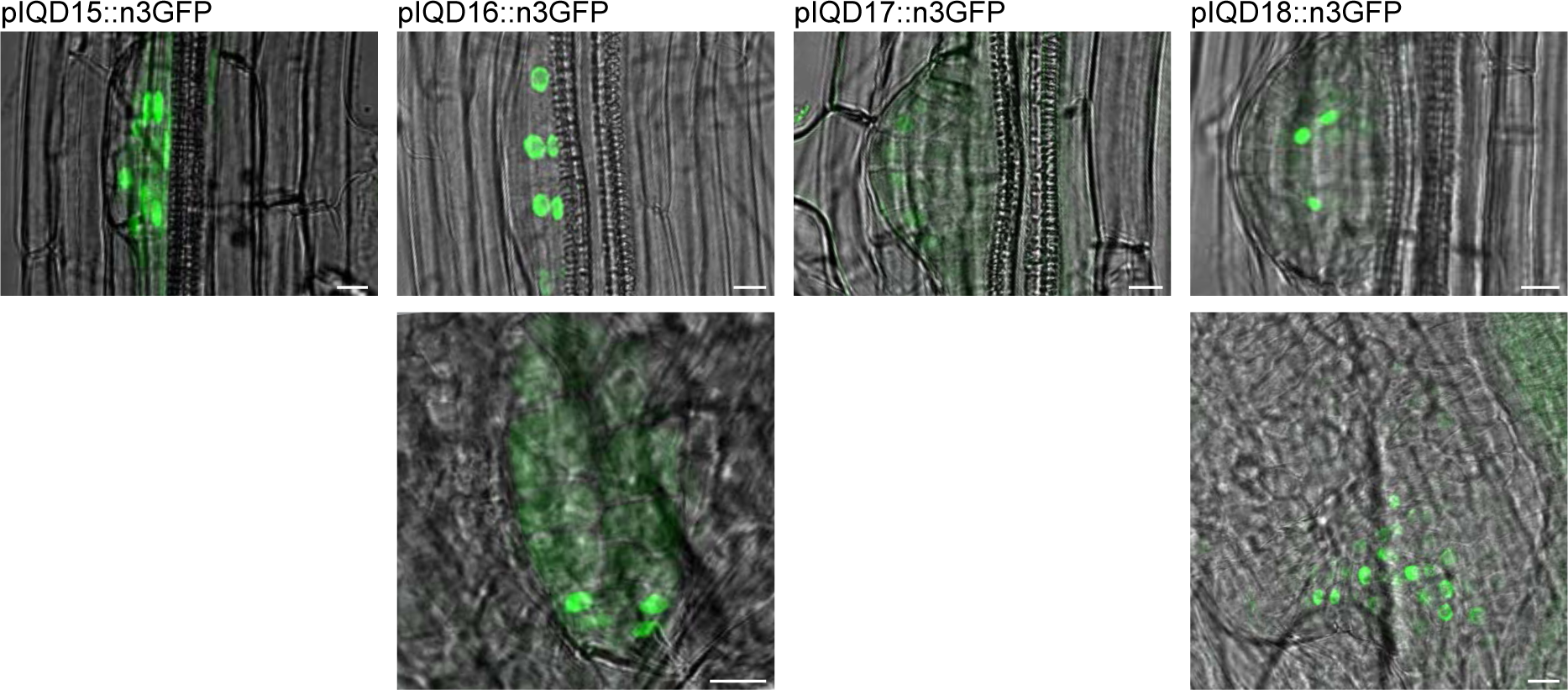
AtIQD expression in lateral root (top) and leaf primordia (bottom), reveals strong expression of AtIQD15, −16 and −18 in young lateral root primordia and weak expression of AtIQD17. AtIQD16 and −18 expression could also be observed in developing leaf primordia of 6-day-old seedlings. Measuring bar = 10 μM.

**Figure S3:**
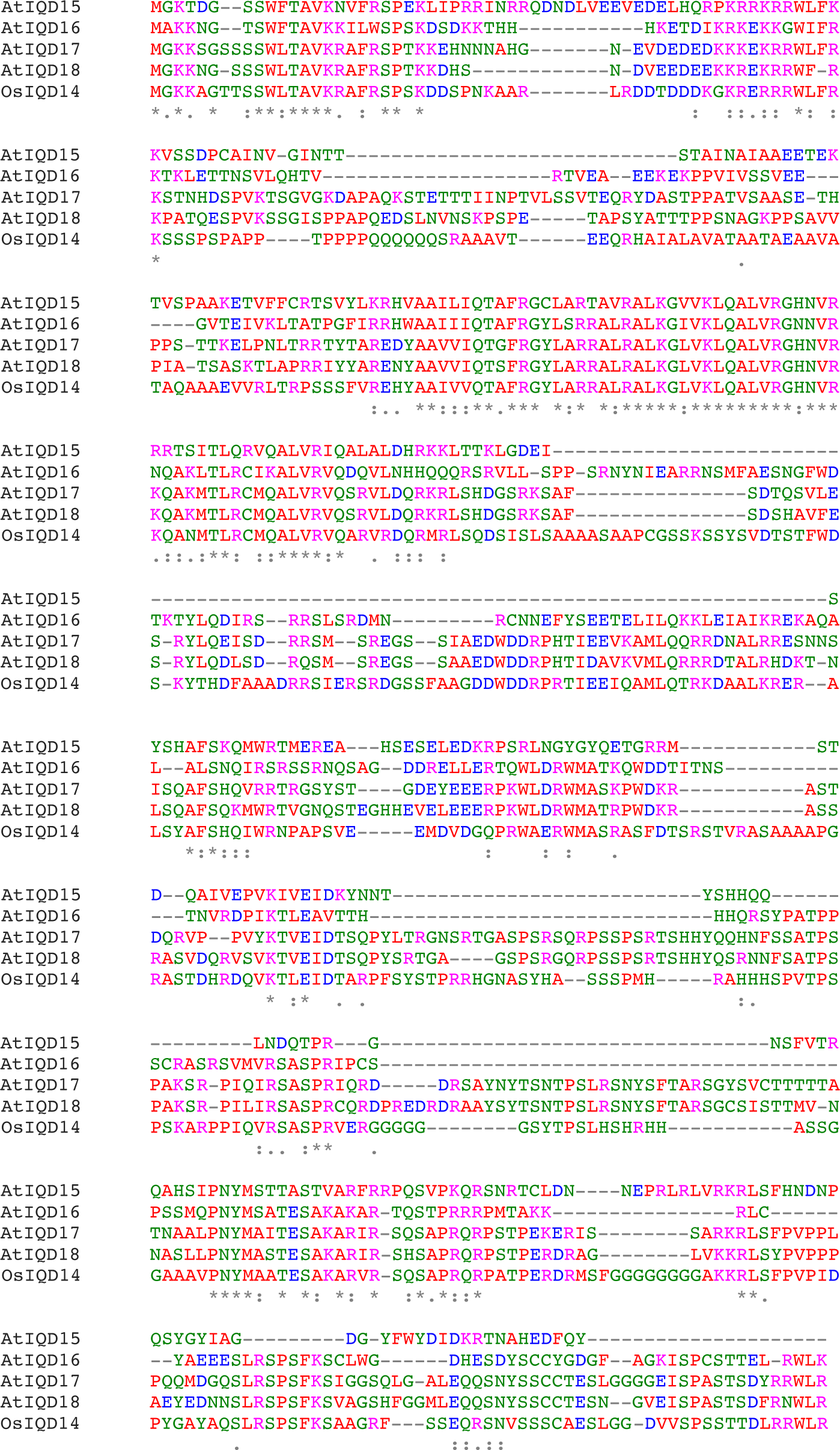
Multiple sequence alignment of AtIQD15-18 and OsIQD14 protein sequences. Conserved amino acids are indicated by semicolons and asterisks indicate highly conserved amino acids.

**Figure S4:**
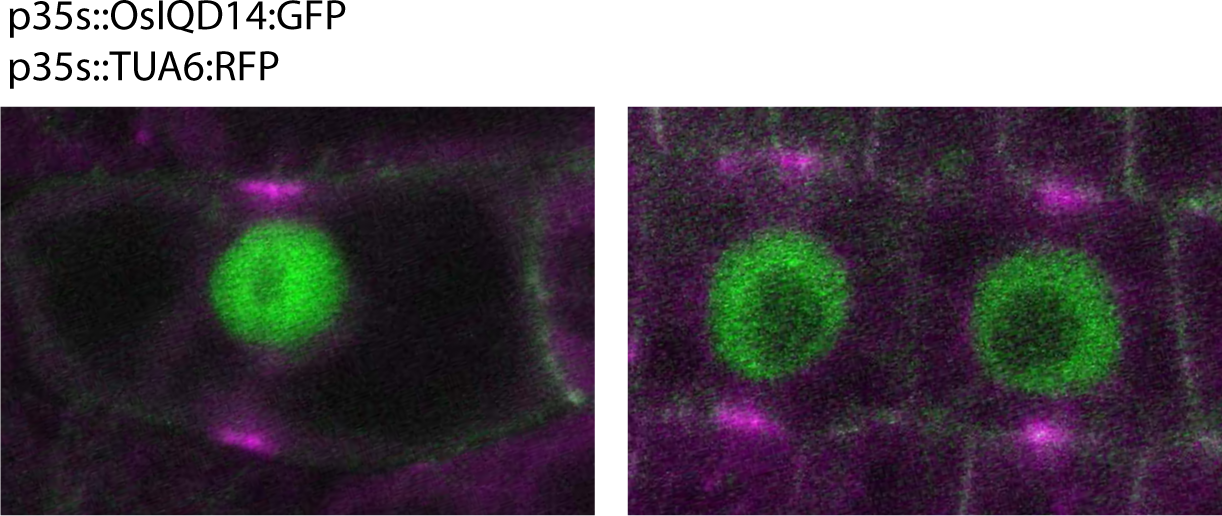
OsIQD14:GFP and TUA6:RFP protein localization in Arabidopsis root cells. Preprophase band is marked by TUA6:RFP (magenta), note GFP signal is excluded from RFP signal.

**Figure S5:**
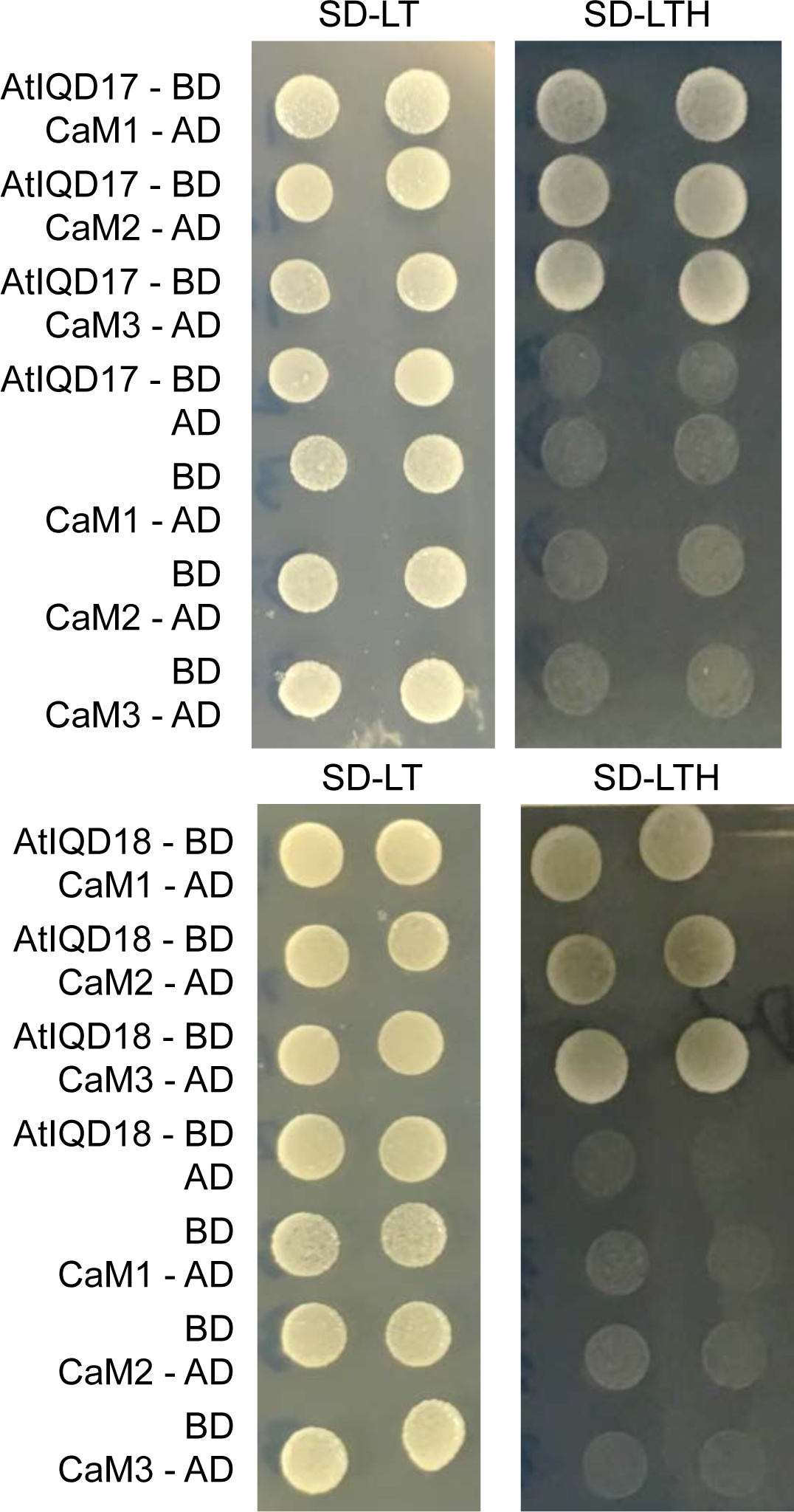
Yeast-Two-Hybrid assays showing yeast growth on selection media (SD-LTH), indicating direct interaction of AtIQD17 and AtIQD18 with CaM1-3.

**Figure S6:**
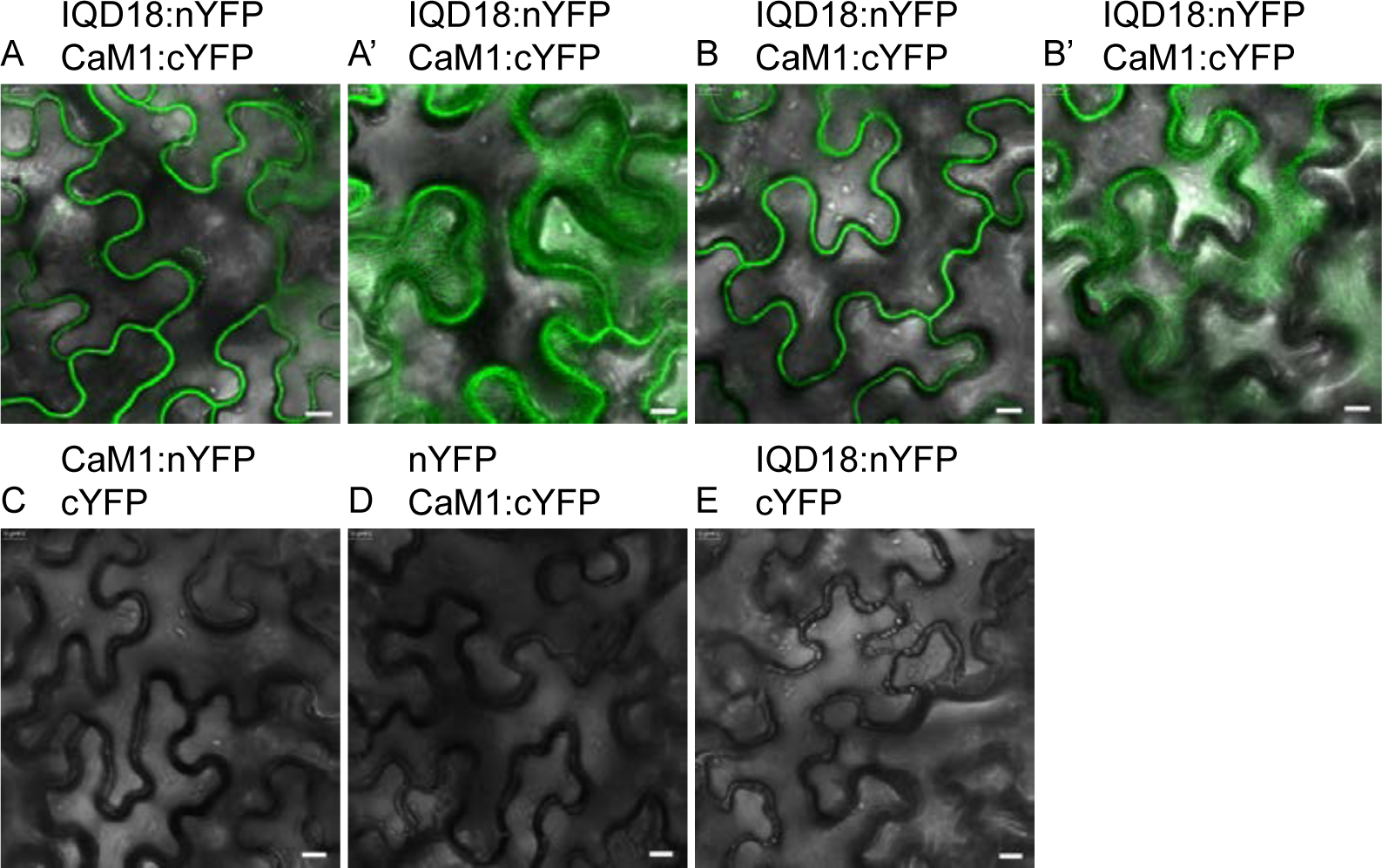
Bimolecular Fluorescence Complementation assays showing direct interaction of AtIQD18 and CaM1 in tobacco leaf cells. A’ and B’ show surface view of pavement cells, revealing binding of AtIQD18-CaM1 complex at cortical microtubules. Measuring bar = 10 μM.

**Figure S7:**
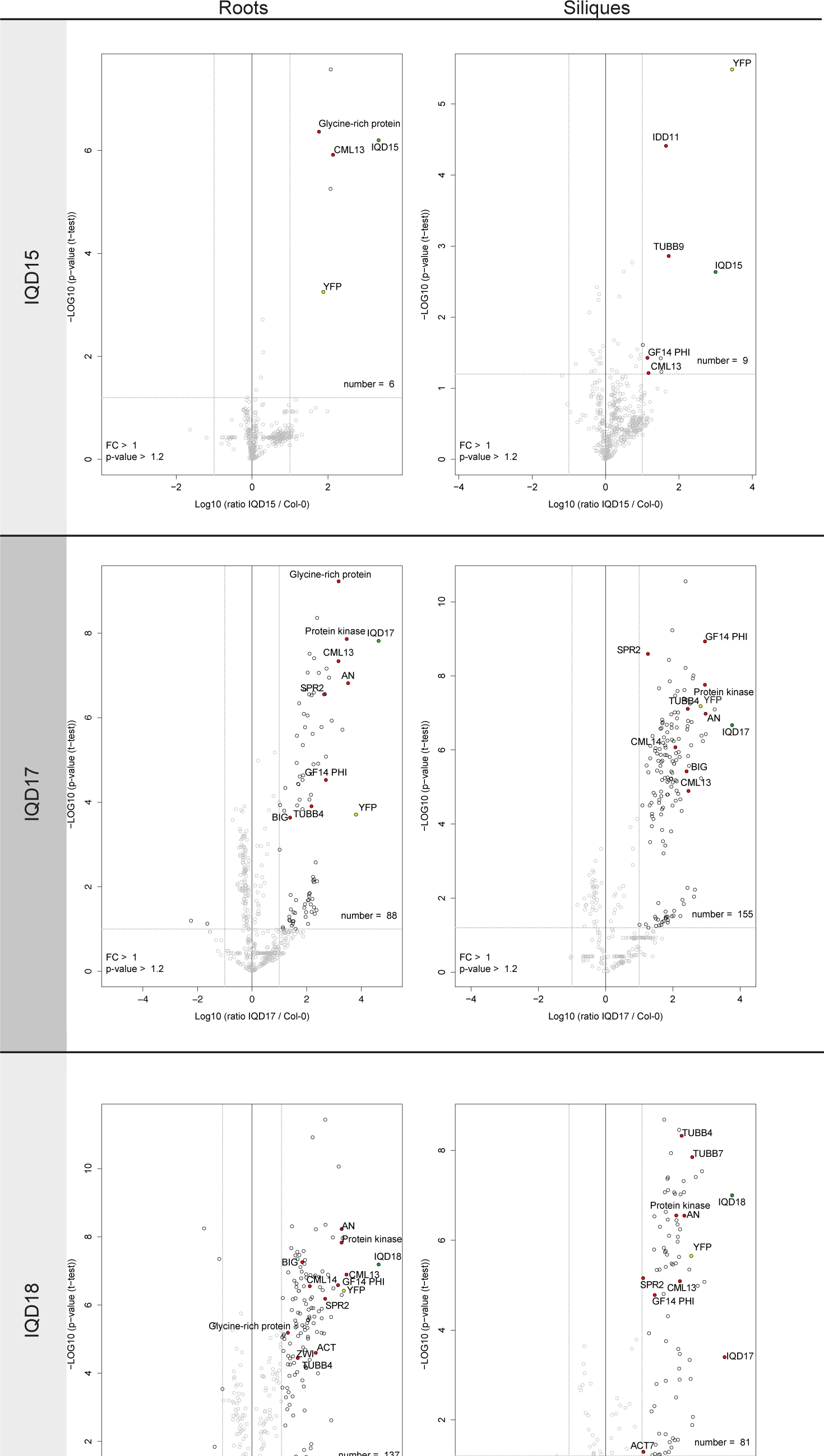
Volcano plots showing significantly enriched proteins in AtIQD15, −17, and −18 IP-MS/MS experiments on root and silique tissues. YFP protein is indicated as a yellow dot and bait protein as green dot. Non-significant and low-ratio proteins are indicated in grey.

**Figure S8:**
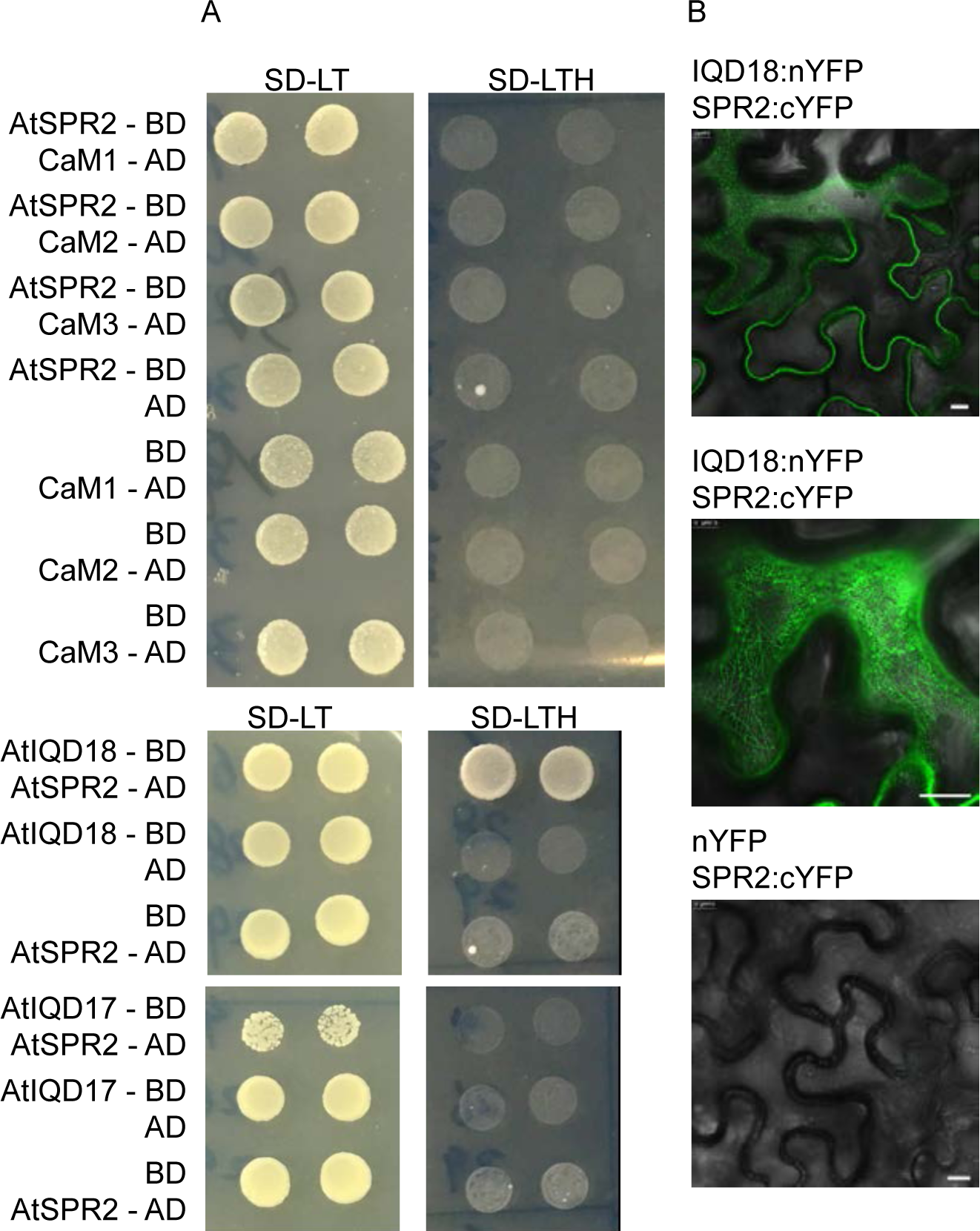
AtIQD18 and SPIRAL2 interact directly at the MT. (A) Yeast-Two-Hybrid assays showing yeast growth on selection media (SD-LTH), indicating direct interaction between AtIQD18 and SPR2, but not between SPR2 and CaM1-3 or AtIQD17. (B) Bimolecular Fluorescence Complementation assays showing direct binding between AtIQD18 and SPR2. Middle panel shows surface vies of pavement cell, revealing binding of AtIQD18-SPR2 complex at cortical microtubules. Measuring bar = 10 μM.

**Figure S9:**
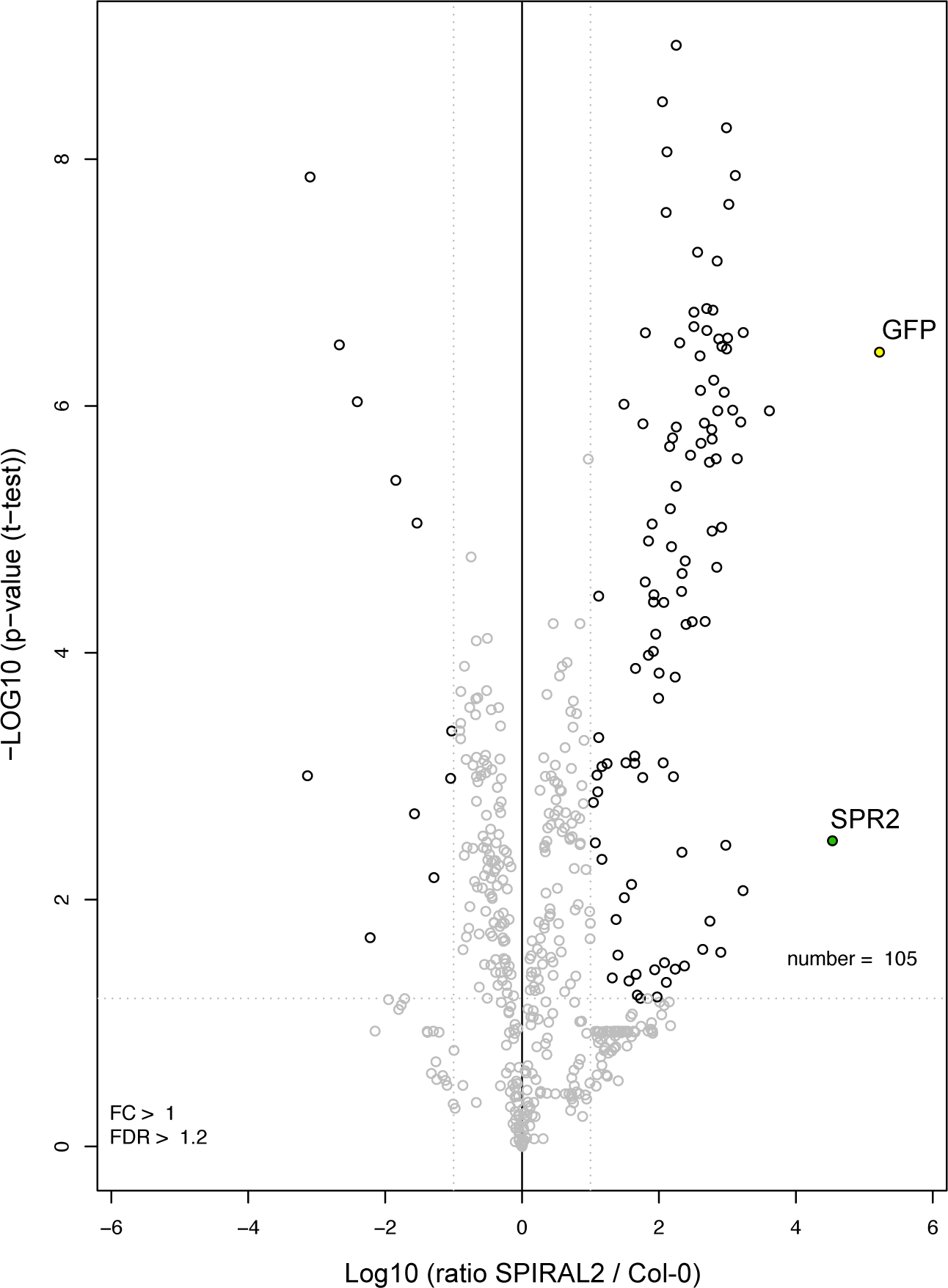
Volcano plot showing significantly enriched proteins in SPR2 IP-MS/MS experiment on whole seedlings. YFP protein is indicated as a yellow dot and bait protein as green dot. Non-significant and low-ratio proteins are indicated in grey.

**Figure S10:**
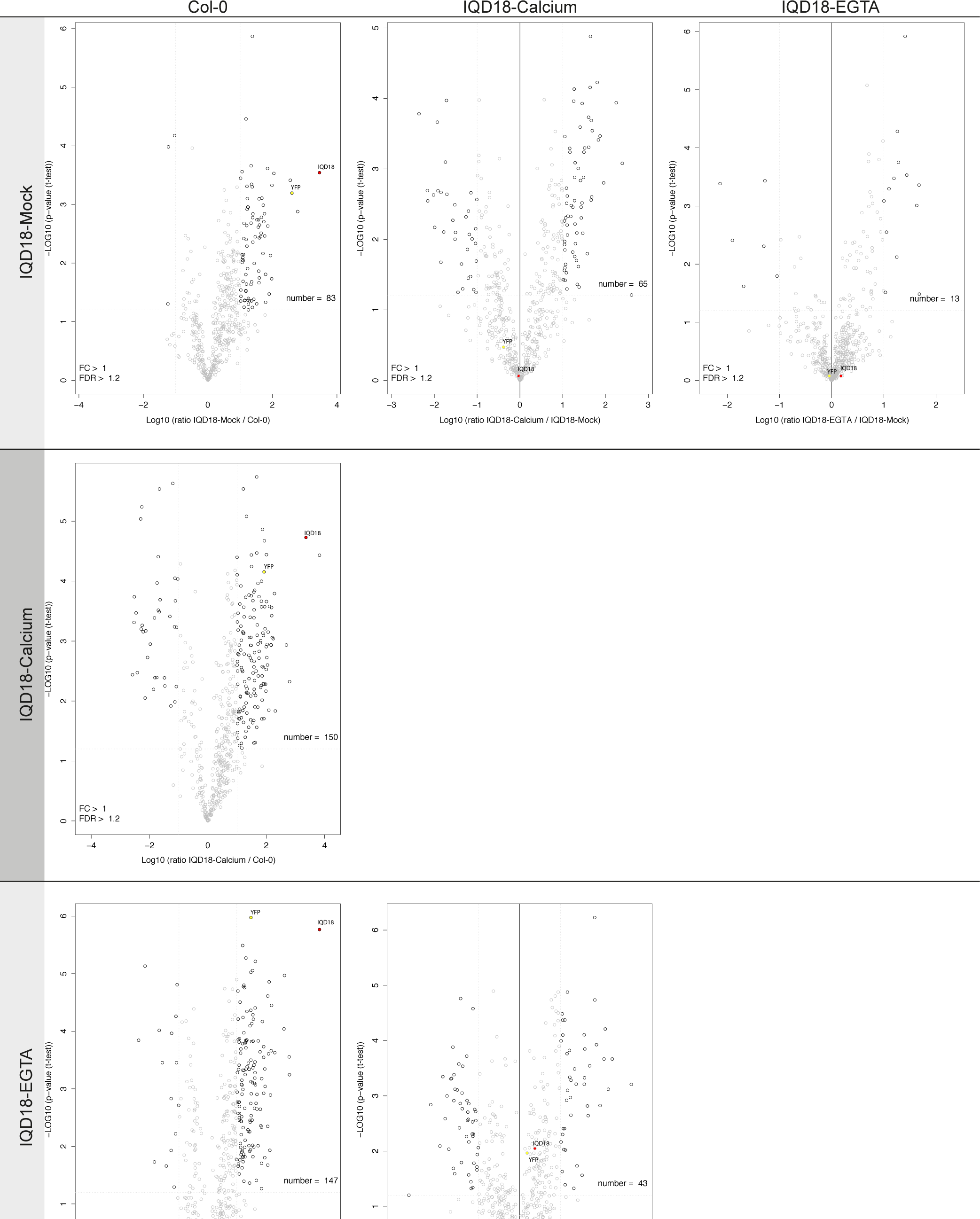
Volcano plots showing significantly enriched proteins in AtIQD18 IP-MS/MS experiments after mock, calcium and EGTA treatments. YFP protein is indicated as a yellow dot and bait protein as green dot. Non-significant and low-ratio proteins are indicated in grey. Comparisons between all treatments are shown.

**Figure S11:**
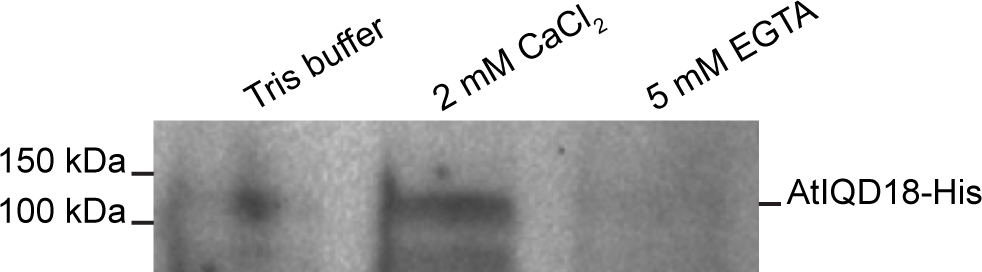
Western blot after pull down experiment using recombinant AtIQD18-His proteins and CaM1-agarose beads, showing increased binding between AtIQD18 and CaM after calcium treatment.

